# Combatting nonidentifiability to infer motor cortex inputs yields similar encoding of initial and corrective movements

**DOI:** 10.1101/2021.10.18.464704

**Authors:** Peter J. Malonis, Ankit Vishnubhotla, Nicholas G. Hatsopoulos, Jason N. MacLean, Matthew T. Kaufman

**Author notes:** Equal contributions (Peter J. Malonis, Ankit Vishnubhotla). Equal contributions (Jason N. MacLean, Matthew T. Kaufman). **Author contributions:** MK and PM conceived the project. PM performed the original analyses of the monkey data and built the simulations. AV analyzed the simulations and the monkey data with ensembles. JM and MK supervised the project with assistance from NGH. NGH provided the data. PM, AV, JM and MK wrote the paper, and all authors edited the manuscript.

## Abstract

Primary motor cortex (M1) plays a central role in voluntary movement, but how it integrates sensory-driven corrective instructions is unclear. We analyzed population activity recorded from M1 of macaques during a sequential arm movement task with target updates requiring online adjustments to the motor plan. Using Latent Factor Analysis via Dynamical Systems (LFADS), we separated neural activity into two components: intrinsic dynamics and inferred external inputs influencing those dynamics. Inferred input timing was more strongly locked to target appearance than to movement onset, suggesting that variable reaction times reflect interactions between inputs and ongoing dynamics. Inferred inputs were tuned similarly for both initial and corrective movements, suggesting a shared input encoding across visually-instructed and corrective movements that was previously obscured by M1 dynamics. Because input inference can suffer from the challenge of nonidentifiability, where different models fit the data indistinguishably, we used ensembles of models with varied hyperparameters to diagnose when inputs are identifiable or nonidentifiable. In the monkey data, ensembles produced consistently similar results, suggesting that inputs could be meaningfully inferred and that their encoding was not simply a result of model bias. These results highlight the challenges of nonidentifiability and the potential of model ensembles to identify inputs in ongoing dynamics, at least in some cases.

## Introduction

The brain is a heavily interconnected network, with many areas working together to achieve control of behavior. Thus, understanding the role of each area and its influence on others requires knowing the inputs to a given region. Here we consider this problem in the context of motor cortex, whose properties make it potentially well-suited for input inference.

Animal behavior requires the motor system to produce reliable movements with precise structure. While some aspects of motor pattern generation occur in the brainstem and spinal cord ^1^, motor cortex exhibits activity patterns with complex temporal structure ^2,3^ that relates to many aspects of movement ^4–8^. However, it remains unclear to what extent activity reflects intrinsically arising pattern generation ^9,10^ versus external sensory and top-down inputs ^11–13^.

One way to understand how the brain generates movement is through dynamical systems analysis ^12,14–18^. In tasks involving brief, largely open-loop reaching, such models approximate motor cortex as a transiently autonomous system with rotational or near-rotational dynamics ^19–22^. These dynamics appear to be initialized by movement-specific inputs ^19,23^ and triggered by large, condition-invariant signals ^24,25^. Yet in continuous, real-world behaviors, the motor system must incorporate ongoing feedback and reconfigure its goals – necessitating inputs external to motor cortex ^15,26^. This reliance on external signals becomes especially clear at movement initiation ^12,27^ or when correcting errors ^15^.

Latent Factor Analysis via Dynamical Systems (LFADS) ^28^ leverages recurrent neural networks (RNNs) to model neural population activity by combining an autonomous dynamical system with a learned stream of time-varying ‘external’ inputs. This architecture allows LFADS to infer single-trial population trajectories using intrinsic dynamics to describe activity where possible, and resorting to inferred external signals as necessary to explain observed activity. Prior work has shown it is highly effective in its goal of fitting denoised single-trial activity trajectories ^28–30^.

Given the success of LFADS in capturing key features of M1 dynamics, we tested whether LFADS could provide insight into the timing and tuning of inputs to M1 during a Random Target Pursuit task ^31^. In this task, the monkey is presented with a new target whenever the current target is contacted, resulting in frequent errors and corrections. LFADS provided a good fit to the data and revealed distinct, large input transients that were closely locked to target appearance, more so than to the movement itself. Notably, input transients encoded target direction during the initial movement and movement corrections similarly.

However, the extent to which generative RNN models like LFADS can accurately disentangle internal dynamics from external inputs has not been systematically assessed. More specifically, it is possible that different parameter values or models produce indistinguishable outputs (here, neural firing rate estimates), the challenge of nonidentifiability ^32–34^.

To address this, we applied LFADS to a toy model simulation with known ground truth. While trajectory estimates were consistently accurate, inference of the input coding scheme was identifiable in some cases and nonidentifiable in others. Fortunately, identifiability could be assessed and diagnosed by fitting model ensembles and analyzing the consistency of their results. In our simulations, when models in the ensemble agreed, inputs were identifiable. Using such an ensemble, we found that the required inference was achievable here.

## Results

We analyzed Utah array recordings from M1 while three monkeys performed a Random Target Pursuit (RTP) task (Fig. 1A). Movement was constrained to the horizontal plane by a two-link exoskeleton. In each trial of this task the monkey made continuous arm movements from target to target, each of which appeared at a random location on the screen when the previous one was hit with a cursor projected just above the hand. The RTP task therefore required continual updating of motor commands in response to stimuli throughout the trial, evoking continuous movement (Fig. 1B,C).

**Figure 1.**
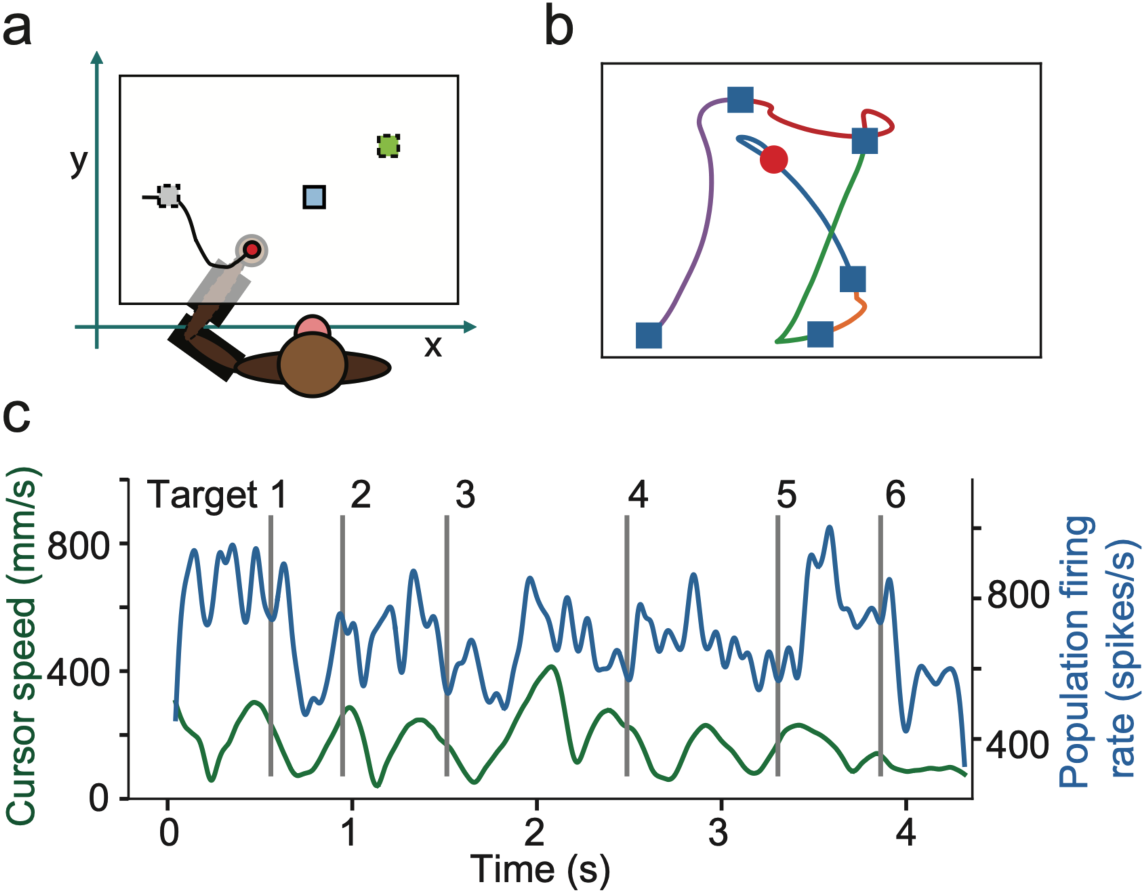
The Random Target Pursuit task elicits continuous movement in response to changing movement goals. **(a)** The monkey performed the random target pursuit task in the horizontal plane using a two-link exoskeletal robot. The monkey moves from a starting position (*gray square*) to the target (*blue square*). Once the target is acquired, a new target immediately appears (*green square*). **(b)** An example hand trajectory from one trial from Monkey RS. Starting position is indicated by the red dot, and the target locations as blue squares. The trajectory for each target is drawn with a separate color. **(c)** The speed profile (*green*) and population firing rate (*blue*) in recorded units for an example trial in Monkey MK. Multiple local minima were present in the speed profile for some targets, indicating submovements / corrections.

### A dynamics-based fit to M1 activity required time-varying inputs

We used LFADS, a machine learning method based on RNNs, to model the neural spiking data. This model treats neural data as arising from a dynamical system, initialized at the start of the trial with a seed that is specific to the upcoming activity, and subject to a time-varying set of inputs that arise from outside the dynamical system (Fig. 2A). This model is structured as an autoencoder: neural data are first transformed into the initial state for each trial, then run through a dynamical system to generate the state of the system over time (conceptually like the factors that would result from Principal Component Analysis), and finally these factors are related back to the same neural activity via a Generalized Linear Model. The non-dynamical inputs form a side loop (Fig. 2A, red path), where both the initial state and errors of the factors (failures to reconstruct the data) inform a stream of corrective external inputs to the dynamical generator. The denoised factors from this model have been shown to reliably distinguish between different conditions on single trials from the neural data alone; the factors and initial states accurately predict external kinematic variables; and importantly for our purposes, the inferred inputs correctly infer the sign of external, physical inputs when movements were perturbed on some trials ^28^. In the framework of the LFADS model, the presence of inputs at a given time in the trial means that the evolution of population activity cannot be parsimoniously explained by the autonomous dynamics of the “Generator” RNN. Here, we focused on the nature of the inferred inputs – specifically, the inputs required by the dynamical system at each time point to reproduce the recorded neural activity during continuous pursuit of random targets.

**Figure 2.**
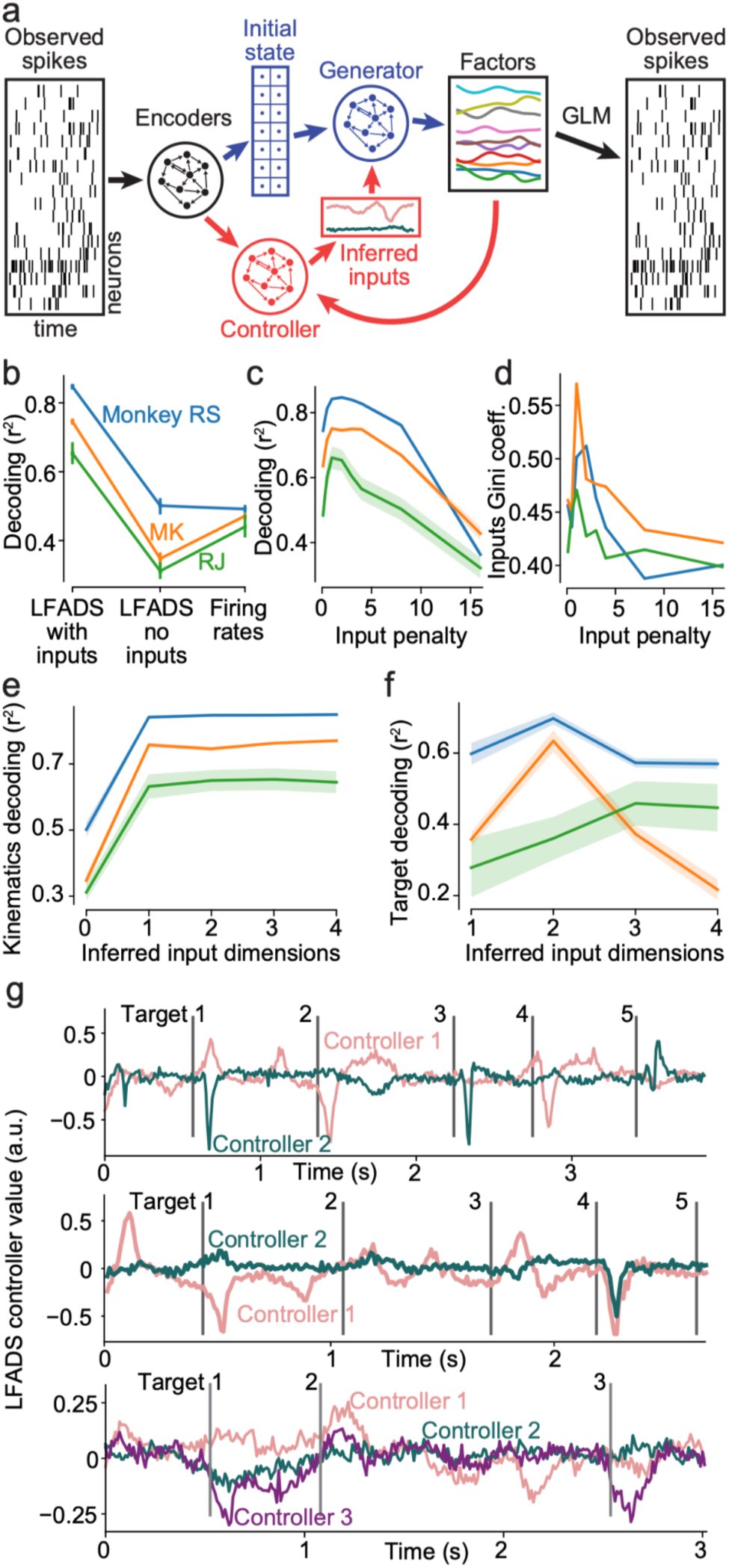
Modeling M1 population activity with Latent Factor Analysis via Dynamical Systems (LFADS). **(a)** Schematic of LFADS approach. For each trial, the spatiotemporal pattern of population activity is reduced to a vector representing an initial state of a dynamical system (*blue*), and a low dimensional time series that represents exogenous input not explained by the autonomous dynamics of that system (*red*). **(b)** Kinematic decoding from LFADS factors for models with and without a controller. Decoding performance using firing rates is shown for reference. Error bars represent standard deviation of performance across cross-validation splits. **(c)** Kinematic decoding performance as a function of LFADS input penalty. **(d)** Gini coefficient of the inferred inputs for models with a range of input penalties. **(e)** Kinematic decoding performance as a function of the dimensionality of inferred inputs used in the LFADS model. **(f)** Target position decoding performances, using the LFADS inferred inputs, as a function of the dimensionality of the inputs used in the model. **(g)** Example inferred input traces for one trial from monkeys RS (*top*), MK (*center*), and RJ (*bottom*).

Because the inference of inputs was the focus here, we took particular care to fit the hyperparameters of the model affecting these external inputs. In general, the same data can be fit as arising from either a more complex dynamical system (higher dimensional or with ‘more nonlinear’ dynamics) receiving less exogenous input, or a simpler dynamical system receiving stronger and more frequent inputs ^35^. For LFADS, whether the system fits a more complex dynamical system or more complex inputs is informed by hyperparameters that are set by the user. If the input-penalizing hyperparameters are chosen to be too weak, too much of the activity will be attributed to the inputs, and not enough of the autonomous dynamics will be learned. If the model penalizes external input too strongly, then the model will not capture updates to the motor plan due to the changing state of the task and may learn overly-complicated dynamics with the initial state distinguishing trajectories that only diverge at late timesteps.

To first evaluate the importance of external inputs to model performance, we measured how well the ongoing movement kinematics could be linearly decoded from the denoised single-trial firing rates, when the model was fit with different dimensionalities and penalties for the external inputs ^28,29^. Including external inputs resulted in better decoding of the movement kinematics than excluding them (Fig. 2B; r^2^ = 0.75 ± 0.08 with inputs, mean ± s.d. across monkeys; r^2^ = 0.39 ± 0.08 without), and decoding from the LFADS single-trial rates was substantially more accurate than using the firing rates directly (r^2^ = 0.46 from smoothed firing rates), demonstrating that LFADS indeed recovered single trial trajectories accurately. Kinematic decoding performance peaked at similar penalty levels and input dimensionality for all monkeys (Fig. 2C,E,F). For further analysis we used the optimal dimensionality for each monkey (2 or 3), and the same penalty level (2.0) for all monkeys.

### Inferred inputs were sparse and locked to target presentation

In principle, external inputs could be continuous or pulsatile, and changes in the inputs could occur sparsely or frequently. However, LFADS penalizes the input magnitude at all time points, and we therefore expected LFADS to remove sustained features in the inputs – which are easy for the autonomous generator to incorporate – much like a high-pass filter. Indeed, the fits produced sparse input values, indicated by high Gini coefficients ^36^ (Fig. 2D). Interestingly, especially given that we used an intrinsically de-sparsifying L2 penalty on the inputs, sparsity was maximal at penalty values close to those at which the denoised trajectories yielded the best kinematic decoding. We therefore focused our analysis on the inputs’ sparse larger values, which allowed us to evaluate how inputs brought new information into the system.

With this optimized model, inferred inputs maintained a relatively low and stable baseline with excursions that were sparse, large, brief, and tended to occur immediately following the presentation of a new target (Fig. 2G). Averaging the magnitude of inferred inputs across target presentations revealed a peak at ∼50 ms after target appearance for all three monkeys (Fig. 3A). The timing of the inferred input peak coincides with previous reports of the latency from visual movement cues to the start of changes in M1 activity ^37,38^. To examine how reliable the peak timing was on individual trials, we computed the latency from the target onset to the first inferred input above a threshold. As the threshold was increased, the peaks became more sharply peaked around 50 ms following target presentation, indicating that the largest transients were the most consistently timed (Supp. Fig. 1A), and that these inputs were larger than at other times in the trial (Supp. Fig. 1C).

**Figure 3.**
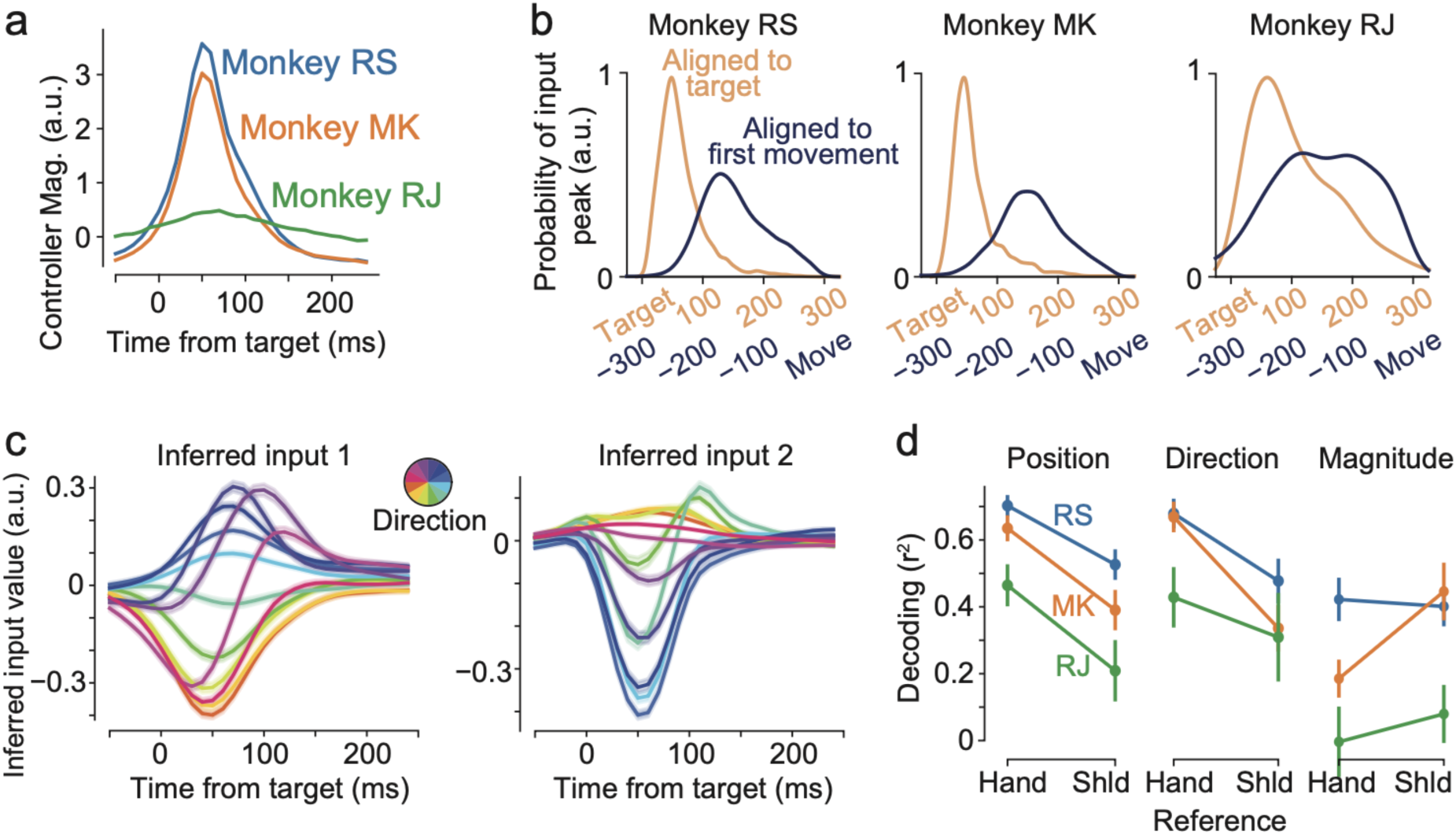
LFADS inputs exhibit transients that are locked to the appearance of new targets and prior to corrective movements, and predict the location of the next target. **(a)** Average controller magnitude relative to target presentation. SEMs are largely hidden by the traces. **(b)** Distribution of the timing of maximum inferred input magnitude aligned to target presentation (*orange*) and movement (*dark blue*). **(c)** Average of the inferred input for different directions of the target relative to the hand. Shading represents ± SEM. Monkey RS. **(d)** Decoding performance on held-out data of target position (*left*), target direction (*center*) and target distance (*right*) relative to both the hand and shoulder. Error bars represent standard deviation of performance across cross-validation splits. Shld, shoulder.

To better evaluate the specific timing of the input transients, we examined whether the timing of the transients was more consistently related to the timing of target presentation or to the timing of movement initiation towards the target. The start of the initial movement was identified by examining the speed of the hand following target presentation and selecting the appropriate speed local minimum (see Methods). For each input peak as defined above, we then examined the latencies to both the target presentation and the initial movement (Fig. 3B). The distribution of latencies relative to the target presentation had both a smaller variance and smaller width at half-height compared to the latencies relative to the initial movement (p < 0.001 for both, permutation test, see Methods). This result suggests that the timing of the post-target input transients were more consistently related to target appearance rather than movement onset. This in turn suggests that controller inputs are related to visual input responses, and therefore that the variable latency of responding to a new target may result from the interaction of these inputs with the dynamical system of M1.

### Inferred input transients relate to target direction in hand-centered encoding

Given the strong correspondence between the timing of target appearance and of the inferred inputs, we evaluated the extent to which the values of the inferred inputs predicted the location of the target. To do so we made a sort of tuning curve by binning targets according to the direction from hand to target, then averaging the inputs for each bin to get a trace over time for each direction (Fig. 3C). Despite the continuous nature of the movement this approach revealed highly distinct inputs for different upcoming movements.

We then decoded target location to verify that the inputs were meaningful, particularly around the time of target presentation (Methods). To do so, we compared the cross-validated decoding performance using either inferred inputs or M1 firing rates in different windows of time relative to the target presentation, with window sizes equal to 250 ms (Supp. Fig. 1B). M1 firing rates yielded a stable prediction of the target position throughout the full duration of the trial, consistent with M1’s known correlations with movement across time ^7,39^. In contrast, using inferred inputs, decoding performance rapidly decreased when the start of the window of input used was >50 ms after the target presentation. This contrast suggests that inputs in the model only briefly encode the movement, but they persistently alter the state of the dynamical system, such that the encoding is sustained in the evolution of M1 activity for the remainder of the movement.

Areas upstream of M1 in the fronto-parietal reach network have been shown to more strongly represent the difference vector from the hand to the target than the absolute position of the target relative to the torso ^40,41^. We therefore hypothesized that similar encoding might be present in the inferred inputs. We evaluated the performance of a nonlinear decoder predicting target position depending on whether the hand or shoulder was used as the origin for each target. Consistent with previous reports, the encoding of the targets by the inferred inputs was stronger relative to the hand than relative to the shoulder (Fig. 3D *left*; r^2^ = 0.70 vs. 0.53 for Monkey RS, 0.63 vs. 0.39 for Monkey MK, and 0.46 vs. 0.21 for Monkey RJ; p < 0.001 for each monkey, corrected t-test, see Methods).

We also trained a decoder to predict the direction and magnitude of the vector pointing from the point of reference (hand or shoulder) to the target. The performance and difference between the hand-centric and shoulder-centric models was similar for the direction decoder (Fig. 3D, *center*). The magnitude decoder exhibited weaker performance in general (Fig. 3D, *right*), with no consistent differences between hand vs. shoulder origin. This indicates that the difference in decoding performance between the hand- and shoulder-centric decoders is largely due to the more accurate representation of target direction by the inferred inputs.

### Corrective movements were also preceded by transient inputs

Many large inputs were time-locked to target presentation, but large inferred input transients were also observed outside of these time windows (Fig. 2G). We hypothesized that these transients might correspond to corrective submovements, which are common in the RTP task (Fig. 4A).

**Figure 4.**
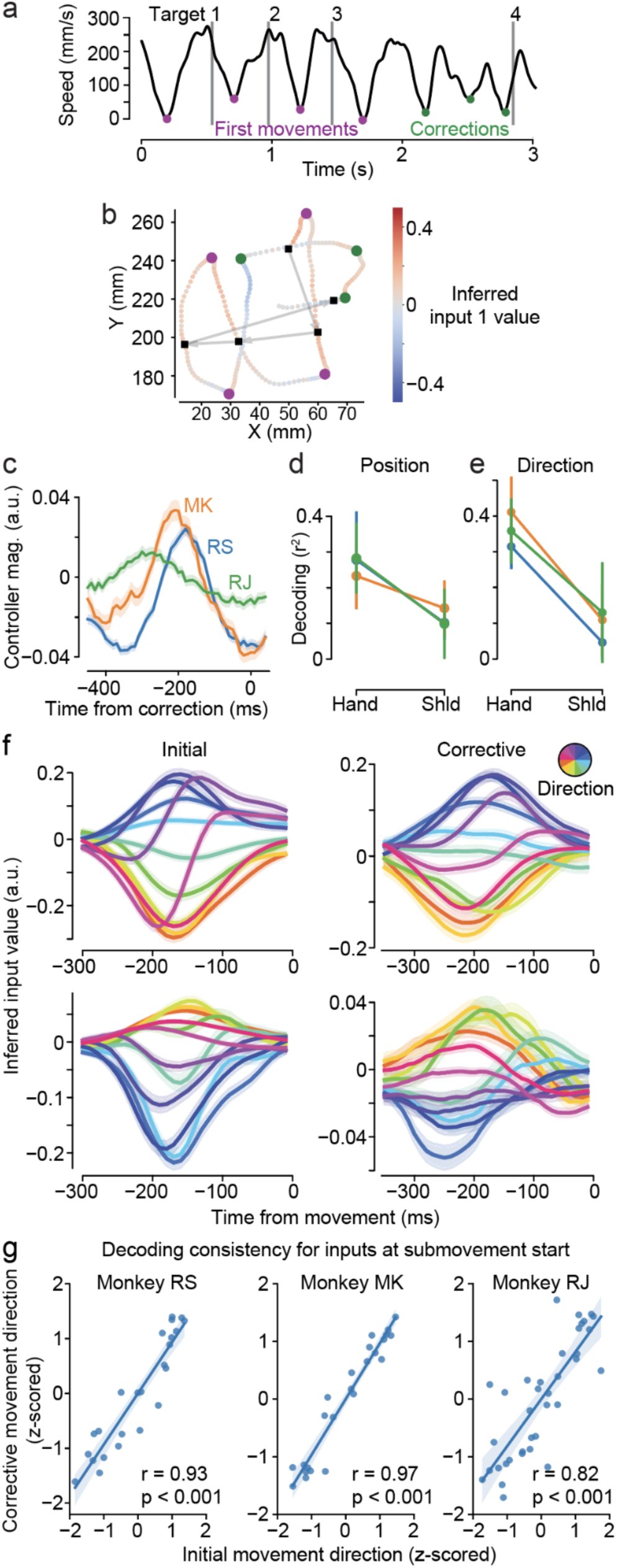
LFADS inferred inputs also predict corrective movements. **(a)** Speed of the hand across an example trial for monkey RJ. Target presentation times, gray lines; initial movements (*purple dots*) and corrections (*green dots*) identified from local speed minima. **(b)** Movement trajectory and inferred input value 1 for the same trial as in **(a)**. Black squares, target location or initial position of hand at start of trial. Arrows, sequence that targets were shown. Dots, position of hand at one LFADS sample (100 Hz). The colors of the pale dot trail represent value of inferred input 1 at that time point. Initial movements and corrections shown as in **a**. **(c)** Average controller magnitude relative to corrective submovement onset. Shaded areas are SEMs. **(d)** Cross-validated decoding of target position and **(e)** direction for corrective movements. **(f)** Average of inferred inputs for monkey RS for different directions for initial movements (*left*) and corrections (*right*). Input 1 shown on top, input 2 on bottom. **(g)** Correlation between mean controller value in window from 350 to 50 ms before the movement for initial movements vs. corrective movements. Each point represents the mean for one direction of movement for one input, z-scored using the mean and s.d. for that input.

In example trials, we found that bends in the hand trajectory were associated with transients in the inferred inputs (Fig. 4B). To quantify this, we identified submovements by appropriate local minima in the speed of the hand (Fig. 4A; see Methods). We defined corrective submovements as all submovements that occurred after the initial submovement that followed the presentation of a new target. We then asked whether the time windows around corrective submovement onsets were associated with transients in the inferred inputs. For all three monkeys, we found a peak in the inferred input magnitude at the start of corrective submovements, 180 ms to 300 ms before the speed minimum (RS: 180 ms; RJ: 300 ms; MK: 210 ms) (Fig. 4C). As we found for initial movements, the transients in the 200 ms period prior to corrective submovements were statistically different from random time windows assessed via the area under the ROC curves (Supp. Fig. 2A, *bottom*). This time window therefore corresponds to the period relative to the initial movement where the inputs are particularly active (Supp. Fig. 2A, *top*).

### Inferred inputs similarly encode both initial movement and corrective submovements

To understand the information content of the inferred inputs for corrective submovements, we trained a decoder to predict the position of the targets from the inferred inputs around corrective movements. This decoder explained the 23-28% of the variance in the hand-relative target position (Fig. 4D), and 31-41% of the target direction in held-out data (Fig. 4E). As with the post-target-presentation inputs, the prediction was much better for target direction when assessed relative to the hand than relative to the shoulder (r^2^ = 0.31 vs. 0.05 for Monkey RS, 0.41 vs. 0.11 for Monkey MK, and 0.36 vs. 0.13 for Monkey RJ, p < 0.001 for each monkey, corrected t-test), and the magnitude decoder again had weaker performance and did not exhibit differences between the hand- and shoulder-centric direction decoders (Supp. Fig. 2B).

Given that the inferred inputs to M1 exhibited transients both following initial target presentation and around the time of corrective submovements, and that the direction of the upcoming movement could be decoded in both cases, we measured the similarity of these two representations. As in Figure 3C, we binned the target directions and computed the average input for each direction (Fig. 4F). Averaging the inferred input for each direction in a window from -350 to -50 ms relative to the start of movement, we produced tuning curves for inferred inputs at target presentation and at corrective submovement onset. The two representations were highly similar (Fig. 4G), with an overall correlation of 0.89 (95% CI [0.83, 0.93]; monkey RS: 0.93, 95% CI [0.84, 0.97]; MK: 0.97, [0.93, 0.98]; RJ: 0.82 [0.67, 0.9]).

This result was intriguing because in general the target representations in M1 are complex, evolving over the course of a trial ^2,42^. Accordingly, we found that the similarity in input encoding was not simply due to similarity in firing rates. We repeated the comparison of tuning curves using the first 5 principal components of the M1 firing rates instead of inferred inputs (Supp. Fig. 2C) and found that the correlation in tuning between the initial movement period and the corrective submovements was much lower, at 0.46 (95% CI [0.34, 0.57]; Monkey RS: 0.53, [0.32, 0.69]; MK: 0.26, [0.01, 0.48]; RJ: 0.57 [0.37, 0.72]). Similarly, tuning consistency was lower when comparing inputs at the time of initial movement and at peak hand speed (Supp. Fig. 2D) than when comparing initial and corrective movements (p < 0.001, Z-test with Fisher’s Z-transformation; Monkey RS, 0.56 vs 0.93, p < 0.001; MK: 0.81 vs 0.97, p = 0.002; RJ: 0.44 vs 0.82, p = 0.005). This argues that the tuning of inputs is more consistent between initial and corrective movements than the tuning of the firing rates themselves or with inputs at other potentially relevant times in the movement.

### Model ensembles diagnose when inputs are identifiable and enable inference

To better understand how hyperparameter choices influence input inference in LFADS, we fit LFADS to activity from a toy model: a continuous-rate RNN simulation controlling a two-link arm executing the random target pursuit task (Fig. 5A; Methods). Specifically, we tested whether and when the inferred inputs from LFADS matched the ground-truth timing and coordinate frame of inputs to the network. We simulated two input timing schemes: one in which the input coordinates were transiently injected into the network, and another in which the inputs were sustained (Fig. 5A). For each timing scheme, the input coordinates were provided relative to either a shoulder or a hand-centered coordinate frame. We then fit LFADS with a variety of hyperparameter choices encouraging it to identify more transient or more sustained inputs, then tested whether the correct coordinate frame could be reliably recovered from the inferred inputs by our panel of decoders.

**Figure 5.**
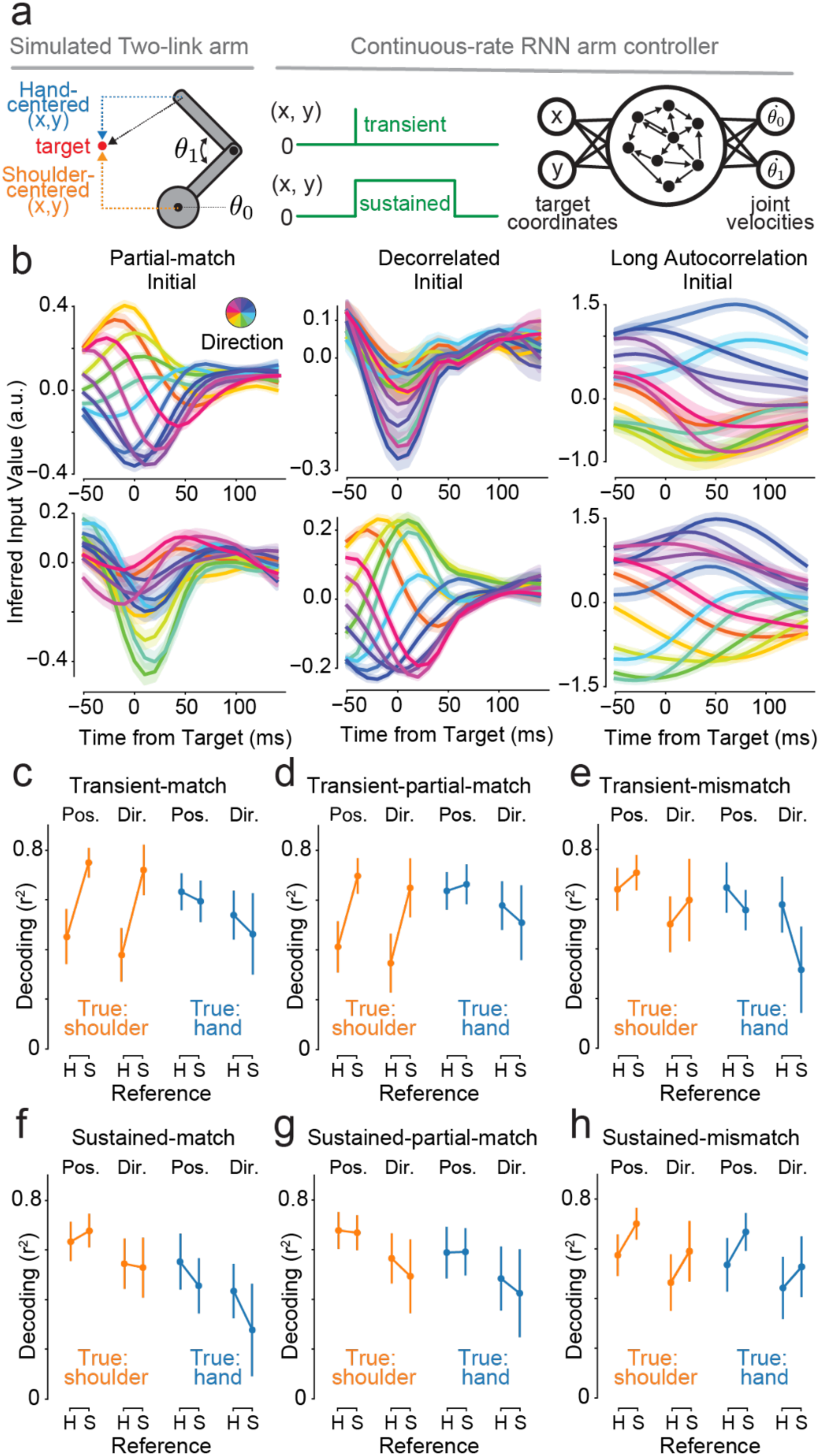
Simulations allowed testing how well LFADS inferred inputs. **(a)** RNNs controlled joint angle velocities of a 2-link simulated arm, guided to targets specified in one or another coordinate frame. **(b)** Inferred inputs as in Figure 3c for a hand-centered, transient simulation and three sets of LFADS hyperparameter values. Inferred inputs for initial movements shown. Sustained simulations produced similar results. **(c)** With transient simulation inputs and matching hyperparameters in LFADS, input inference worked well. **(d-e)** With transient simulation inputs and partially matching or mismatching LFADS hyperparameters, input inference was sometimes correct and sometimes inconclusive. **(f-h)** With sustained simulation inputs, LFADS input inference was generally inconclusive.

A prerequisite for accurate inference was agreement across the ensemble of models: almost by definition, only when multiple models converged on the same encoding were the inputs potentially identifiable. When models in the ensemble disagreed, this meant that inputs were nonidentifiable. For potentially identifiable cases, it was then essential to check that the consensus solution agreed with ground truth.

First, we verified that the hyperparameters directly affected the time course of the inferred inputs. We tested hyperparameters that either encouraged a short time course and decorrelation of the inputs over time (“match” for transient simulations and “mismatch” for sustained simulations), that only encouraged a short time course but not decorrelation (“partial match”), or that encouraged a long time course without decorrelation over time (“match” for sustained simulations and “mismatch” for transient simulations; Methods). As expected, settings favoring transient inputs produced briefer time courses, whereas hyperparameter settings favoring sustained inputs produced more sustained time courses (Fig. 5B).

For transient ground-truth inputs, ensemble agreement of inferred input encoding was always achieved, regardless of whether hyperparameters favored transient or sustained inputs. In these cases, decoding from the inferred inputs recovered the correct coordinate frame and achieved statistical significance (Fig. 5C–E), though in one case the significance was marginal. The potential to accurately recover transient inputs suggests that, for this simulation, time course and hyperparameter range, the encoding scheme of the inferred inputs was identifiable.

In contrast, for sustained ground-truth inputs, ensemble agreement was never achieved. Even when hyperparameters were set to encourage sustained-input solutions, inference of the correct coordinate frame was inconsistent at best (Fig. 5F-H). In the other hyperparameter set cases, coordinate-frame decoding shifted to being either entirely inconclusive or disagreeing. Sustained inputs were therefore nonidentifiable in this setting.

Taken together, these results show that LFADS can usually identify transient but cannot identify sustained inputs in this toy system. More broadly, input identifiability cannot be predicted *a priori* from hyperparameters or model architecture alone. Instead, it must be established empirically: when an ensemble of models converges on the same solution, inputs are likely identifiable; when they do not, inference remains nonidentifiable.

### Input inference was identifiable in the monkey data for two of three animals

Building on the diagnostic role of model ensembles, we repeated our inference on the monkey data using varied input-influencing hyperparameters. In two of the three animals (RS and MK), decoding converged on clear and consistent conclusions for initial movements (Fig. 6A-C). For one monkey (RJ), ensemble results were generally in agreement but some hyperparameter combinations produced weak results. This suggests that inputs for monkey RJ were marginally identifiable or nonidentifiable. For the two identifiable datasets, corrective movement encoding was highly correlated with initial movement encoding in every case (Fig. 6D), supporting the conclusion that input coding was nearly identical for initial and corrective movements. For the marginally identifiable monkey, this correlation was much weaker. Thus, the diagnostic test of ensemble agreement suggests that inputs were identifiable in two of the three datasets, enabling inference of input coding in the monkey data.

**Figure 6.**
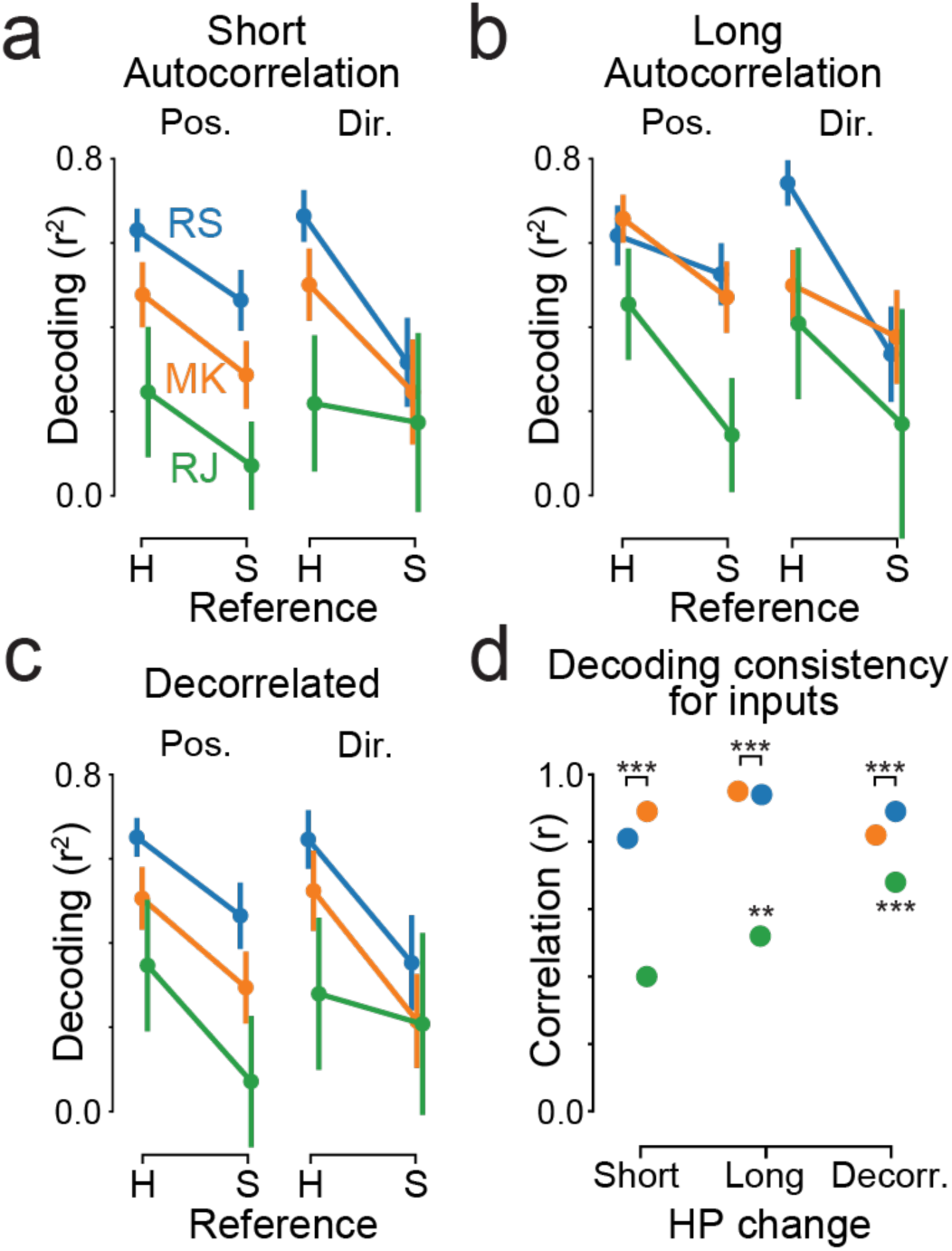
LFADS ensembles agreed in the monkey data. **(a-c)** For monkeys RS and MK, LFADS models trained with hyperparameters encouraging short (**a**) or long (**b**) autocorrelations, or using decorrelated inputs (**c**), produced consistently hand-centered encoding for the inputs preceding initial movements. For monkey RJ, the ensemble sometimes produced weak results, indicating that those data were only marginally identifiable or nonidentifiable for input coding. **(d)** Correlations between input coding for initial and corrective movements (as in Fig. 4g) were high for the two monkeys for whom inputs were identifiable. Results were inconclusive for the nonidentifiable monkey, RJ.

## Discussion

We used a variational RNN-based autoencoder, LFADS, to investigate the structure of population activity in M1 during a continuous reaching task in which targets were frequently updated. This task, by design, required the integration of ongoing sensory feedback and frequent movement corrections, providing an ideal setting to test whether motor cortical activity could be decomposed into internal dynamics and external inputs. LFADS, built to infer single-trial latent trajectories by modeling population activity as a transiently-autonomous dynamical system intermittently perturbed by time-varying inputs, fit the data well and revealed prominent input transients following target onset and prior to corrective movements. Moreover, the format of the inferred inputs was strongly similar between initial and corrective movements. These observations align with prior reports of externally driven modulation in motor cortex during feedback-driven behavior^12,43^.

To validate the input inference process, we used toy network simulations with known ground truth. While LFADS accurately and reliably recovered single-trial population trajectories, its ability to distinguish between activity driven by internal dynamics and external inputs is less directly constrained by the nature of the data and consequently has the potential to be nonidentifiable. That is, in some cases different hyperparameters can produce different solutions with indistinguishable predictions about the observed data. This “undetermination” or “nonidentifiability” make input inference impossible in these cases ^32–34^. Put another way, because a family of models can fit the data similarly well, it is difficult to determine which model most accurately reflects the true balance between internal dynamics and external inputs. Whether a given question asked of a given dataset corresponds to an identifiable property is hard to determine *a priori*.

Encouragingly, our simulations suggested that using ensembles of models could serve as a diagnostic determining whether inputs were identifiable or nonidentifiable. When results of the ensemble generally agree, this suggests the input was identifiable. In our case, even with hyperparameters that were somewhat mismatched to the data, LFADS either correctly identified the coding scheme of the inputs or produced an inconclusive result. We therefore suggest that using an ensemble of LFADS models with a few different choices of hyperparameters can be informative about how well-identifiable the system is, and when the ensemble members agree, the fit is likely to produce accurate information. From our simulations, networks with transient inputs yielded better ensemble consensus and agreement with ground truth than networks with sustained inputs. This could follow from a number of factors, including the architecture of LFADS, but also agrees with previous findings that transient inputs constrain the network architecture to a smaller solution space ^44^.

In the future, our results about input coding could be validated experimentally using calcium imaging, which permits identifying and recording input neurons. With the two-photon imaging variant of LFADS, RADICaL ^45^, inferred inputs could also be compared with biologically-identified ground truth.

Given that we have demonstrated the potential for successful input inference with well-chosen hyperparameters or ensembles, we cautiously interpret our results from the monkey data below. We first expressly note the caveat that we do not know the appropriate values of the hyperparameters, and could not explore the full hyperparameter space due to computational cost. Nevertheless, several factors suggest that our results are reliable. First, our ensemble showed strong agreement, consistent with input identifiability in these data. Second, as internal checks, the inferred input timing consistency, sparsity, presence before corrections, and decodability all peaked for similar hyperparameter values. Third, as an external check, our results for initial movements match results about the coding scheme in parietal cortex ^41,46,47^.

If the inferred inputs are accurate, the similarity of inputs for initial target presentation and corrections would be surprising: firing rates are tuned somewhat differently for delayed and non-delayed movements ^48,49^, and generally exhibit inconsistent tuning at different points in the trial^2,7^. Previous work on corrective movements has proposed at least two distinct processes that produce corrections during arm movements ^26,50^: online modifications to the initial impulse, as studied here, and subsequent discrete position adjustments that bring the end effector to the target. In the latter, discrete-movement case, recent work has argued that firing rates differ in their coding between initial and subsequent corrective movements ^30^. Our findings focus mainly on online correction, but suggest that these processes may be more unified than previously thought. First, the similarity of input tuning for initial and corrective movements suggests that the rest of the brain may not have to make a strong distinction between instructions and corrections; observed differences may come from the interaction of the inputs with a system in a different state. Second, the lack of inputs around the time of max speed does not support previous supposition about the timing of an initial impulse correction. Our results may indicate a greater compartmentalization of motor function than thought, with M1 acting as a movement generator that incorporates movement context, while inputs convey goal information without requiring complex modulation to reflect current movement details ^43^. However, if a strong bias were present in the inference process, it could also lead to this result.

Our findings could also have implications for the understanding of reaction times. It has previously been argued that reaction times reflect variable timing of the movement commands themselves^51,52^, possibly due to sensory variability ^53^ (Osborne et al., 2005) or self-monitoring of readiness ^2^, and commensurate with M1 firing rates being strongly locked to movement onset ^54^. However, here the inferred inputs were most tightly locked to the target presentation itself, not the movement onset. This could suggest that the variable reaction time of the monkey may be at least partly due to an interaction of the input with the current state of the dynamical system, and that depending on this state the system may be faster or slower to initiate. However, it also is possible that this is only true in a continuous-movement task like RTP.

Interestingly, inferred inputs were best described as encoding target position in hand-centric coordinates rather than body-centric, consistent with representations in cortical areas upstream of M1 in the fronto-parietal reach network such as posterior parietal cortex ^46,47,55^, and dorsal premotor cortex ^41^. This result is also consistent with findings that the slow component of M1 and PMd activity is quasi-linearly related to kinematics ^22^, which could reflect inputs that help set up the dynamical system ^23^. Reference frames found across the parieto-frontal reach network are highly heterogeneous ^39^ and this type of description leaves out important aspects of motor responses ^11,13,56,57^. The specifics of coordinate frame findings should be interpreted with caution: this means of analysis presents a binary comparison when the answer may be that neither option is correct. Nevertheless, it is useful in comparing findings with those in parietal cortex in particular, and provides a means for comparing inputs for initial and corrective movements as done here. Thus, to the extent that reference frames capture encoding, our work suggests that inputs to M1 may be closer to hand-centric, both to specify an initial movement and to correct one.

## Methods

### Behavioral task

Three adult male rhesus monkeys (*Macaca mulatta*) were operantly trained to control a cursor in a two-dimensional workspace using a two-link robotic exoskeleton ^58^ (KINARM, Kingston, ON, Canada). The animals sat in a primate chair with their dominant arm in the exoskeleton. Their shoulder joint was abducted nearly 90° and supported by the manipulandum such that all movements were made within the horizontal plane. Direct vision of the limb was precluded by a horizontal projection screen above the monkey’s arm. Visual feedback was available via a visual cursor projected onto the screen. Cartesian coordinates of the visual cursor were determined by digitizing the shoulder and elbow angle at 500 Hz and transforming these variables into a visual cursor position (in centimeters) using the forward kinematic equations for the exoskeleton.

In this configuration, the monkeys performed a random target pursuit (RTP) task. This task required the monkeys to move a cursor (6.7 mm in diameter for monkey RS; MK: 6.7; RJ: 10) to a series of 7 square targets (10 mm on a side for monkey RS; MK: 13 mm; RJ: 20 mm). When the cursor reached the target, the target immediately disappeared and a new one appeared at a random location in the workspace. In Monkey RJ, it was necessary to exclude some trials where the monkey was disengaged in the task for part of the trial. Trials were excluded if the monkey spent more than 500 ms total with a cursor speed less than 1 mm/s.

The raw cursor position was filtered both forwards and backwards with a 3rd order low-pass Butterworth filter with a cutoff frequency of 20 Hz. After computing the velocity via numerical differentiation, the same filter was applied again. Because the algorithm requires trials of equal length, trials were truncated to a fixed length, and trials that were shorter than this length were excluded. The trial length was selected for each data set so as to maximize the total number of time bins across all of the included trials (the time bins per trial times number of trials retained).

### Neural recordings

The neural recordings were collected using a microelectrode Utah array composed of 100 silicon electrodes (1.0 mm electrode length; 400 μm inter-electrode separation). The arrays were implanted in the arm area of the primary motor cortex (M1) of each monkey. During a recording session, signals from up to 96 electrodes were amplified (gain, 5000), bandpass filtered between 0.3 Hz and 7.5 kHz, and recorded digitally (14-bit) at 30 kHz per channel using a Cerebus acquisition system (BlackRock Microsystems, Salt Lake City, UT). Only waveforms (1.6 ms duration) that crossed a threshold were stored and spike-sorted into single units using Offline Sorter (Plexon, Inc., Dallas, TX). For each monkey, a single recording session was included, with 100 neurons, 49 neurons, and 51 neurons from monkeys RS, MK, and RJ, respectively.

### Identification of submovements

Submovements were identified using the speed of the cursor over time. Candidates for the start of submovements were identified as the local minima in the smoothed speed profile. The smoothing was performed using a Savitsky-Golay filter with a window of 101 ms and polynomial order of 2. These candidates had to satisfy two additional criteria to be accepted as the start of a submovement. First, we required that the peak speed of the ensuing movement was at least 25 mm/s greater than the speed at the local minimum. This excluded noise and small movements during periods of hesitancy. Second, we required that the minimum time interval between candidates was 200 ms. This limited the extent to which time windows around submovements overlapped. For candidates within 200 ms, we applied a pairwise selection rule to eliminate one candidate until all candidates were a minimum of 200 ms apart. This selection rule was as follows. For each candidate we computed the speed increase: the next peak speed minus the speed at the minimum. In most cases we selected the candidate with the larger speed increase. However, there were rare cases in which a pair of candidates represented, respectively, the beginning of a submovement and a brief, shallow deceleration interrupting the acceleration phase of the submovement. In these cases, we aimed to define the first candidate as the beginning of the submovement and overrode the “larger increase” rule. These cases were identified if two conditions were both met. First, the dip was shallow: the difference between the preceding speed maximum and the second candidate was <10 mm/s. Second, the speed increases for the two candidates were similar, defined as neither being more than 5 times greater than the other.

Once the submovements were defined, they were identified as initial movements or corrective submovements. The initial movements were defined as the first movement following the target presentation. Corrective submovements were defined as submovements occurring after the initial submovement but before the presentation of the next target.

### LFADS fitting and model evaluation

The spiking data recorded during each trial were modeled using Latent Factor Analysis via Dynamical Systems (LFADS), described in detail in ref. ^28^. This model uses a type of variational autoencoder ^59,60^ to learn a dynamical representation of each trial. The central structure of this network consists of two artificial recurrent neural networks (RNNs), an “encoder” and a “generator.” The generator is trained to produce autonomous dynamics that can be mapped onto the neural activity of a trial. The encoder learns to map the activity of a trial onto a distribution of initial states based on how likely they were to produce the activity on that trial. In addition, a “controller” RNN can be added that learns to compensate for the dynamics that cannot be learned by the autonomous generator. The output of the controller is a distribution of inputs to the generator that are passed on each time step. To obtain the inferred inputs to the neural population for each trial, LFADS samples from the distribution of controller outputs and averages the samples.

Most of the LFADS hyperparameters were adapted from those used for similar recordings in ref. ^28^, with the number of factors outputted from the generator set to 50. The full set of hyperparameters is listed in Table 1. Because we were most interested in the inputs from the controller to the generator in the LFADS model, we most thoroughly explored the hyperparameters directly responsible for determining the structure of the controller outputs (the inferred inputs). These were 1) the penalty on the inferred inputs and 2) the dimensionality of the inferred inputs, (|**u_t_**| in ref. ^28^), including values of zero. The need to systematically explore the influence of hyperparameters here motivated our use of the original LFADS instead of AutoLFADS ^45^, which automatically estimates hyperparameter values.

Models were evaluated by training a linear decoder of movement kinematics from the LFADS factors and evaluating the decoder’s performance (see below). We evaluated a range of inferred input penalties, ranging from values low enough to produce overfitting to values large enough to yield underfitting.

After performing this search we selected an inferred input penalty (= 2.0) that was at or near optimal decoder performance for all monkeys. We fixed the inferred input penalty to this value for the search of dimensionality of the inferred input. In the search for the dimensionality of the inputs, the decoder performance plateaued at 1 input and larger. We therefore selected the input dimensionality for each monkey using the decoding performance for the target (see Fig. 2).

### Decoding of movement kinematics

To evaluate the LFADS models, we decoded movement kinematics from the LFADS factors. An ordinary least-squares linear model was used to map the factors onto the position and velocity of the endpoint (hand), with a 100 ms lag between the LFADS factors and the kinematics. To determine the decoder performance, we performed 5-fold cross-validation, splitting the time bins into random test sets. Performance of the model was evaluated using the held-out variance explained. Performance across the four kinematic variables was averaged to obtain a single measure of performance for each training/test set. For the controls in which kinematics were decoded directly from neural activity, we obtained continuous firing rates by smoothing the raw spike trains with a Gaussian filter with s.d. equal to 25 ms.

### Analysis of input timing

Analyses were performed on the time-varying inferred inputs for each trial. We used all trials in the analysis of the inferred inputs, including both those that were used for training and validation in training the LFADS network. For analysis of inferred input timing, and decoding from inferred inputs below, we excluded the response to the first target on each trial in order to eliminate artifacts resulting from the variable behavior of the monkey before the start of the trial. To determine whether the timing of the input transients was more closely related to the target presentation or to the initial movement (Fig. 4C), we examined the timing of the maximum inferred input magnitude in the period between the presentation of the target and the initial movement. We included any target presentation in this analysis, regardless of how large the maximum input magnitude was. Since the timing of the transients of the individual inputs is more exact when the peaks reach a larger threshold (Fig. 4B), and for some directions there is on average only a small transient (Fig. 5A) which may not be detected, taking the timing of the maximum is conservative for detecting systematic timing precision. But by not setting a threshold for input transients to use, we included all relevant target presentations and thus captured the structure of timing in the model in the most general way.

To compute statistics for Figure 4C, we used a permutation test to evaluate which distribution of latencies had the larger spread. Spread was defined as the full width at half-height (FWHH) of the peak of the distribution. First we subtracted the modal value from each set of latencies to center the peak of both latency distributions on 0. Then we combined the two sets of latencies into a single population that we split in half randomly 1000 times. For each of these random partitions we computed the FWHH, and took the absolute difference between the FWHH of the two random subgroups. These absolute differences constituted the null distribution for the permutation test. We compared the difference in FWHH between the actual samples of the distribution and the null distribution of spreads to obtain a p-value for the hypothesis that the distribution of latencies to the different events had the same spread. Because by definition the maxima we examined were before the initial movement, there was a sharp cutoff on the right side of the distribution of latencies to the initial movement (Fig. 4C). However, this simply means that the FWHH measure is possibly conservative in detecting the greater spread of the initial movement distribution.

We also considered the possibility that the inputs were better aligned to the acceleration phase of the movement. We performed the same analysis, but defining the initial movement time by the time at which the movement reaches half of its peak change in speed. Defining the initial movement in this way did not affect the results or significance levels.

To determine whether windows near target appearances could be distinguished from all other windows, we calculated the area under the receiver operator characteristic (ROC) for each window. To do so, we computed the distribution of input magnitude for a 100 ms long window of interest at a fixed latency to the target. We then compute a “background” distribution of input magnitude in all other 100 ms windows outside this period. We then computed the area under the ROC, which is equal to the probability that the input magnitude in a randomly drawn window at the latency of interest is greater than a random window in the background distribution.

### Decoding from inferred inputs

We decoded target direction from all the samples of inferred input in 250 ms windows around either target presentation or corrective movement. We first split the targets into a training and test set, with each containing 50% of the data. We optimized the decoder for each monkey on the training set. As part of this optimization, we chose between two methods of decoding, support vector regression and random forest regression, using the scikit-learn implementation of both methods ^61^. The optimization procedure included fitting what are generally the most important hyperparameters for these two methods ^62^, as well preprocessing parameters. For the support vector regression, we used a radial basis function and fit the L2 regularization hyperparameter from a range of 0.1 to 1.5. All other parameters were set to the scikit-learn defaults. For the random forest regressor, we fit the minimal samples for each leaf (from the values 1, 5, 10, 15), and the maximum number of features to consider at each split. The number of features (optimized) was either the full number of features, the square root of the number of features, or the base 2 logarithm of the number of features. Two preprocessing parameters were optimized: (1) the time window relative to the reference event to use, and (2) the lag between the event and the hand position when computing the hand-centered direction to the target. The decoder hyperparameters were optimized using 5-fold cross validation on the training set. We then evaluated the selected decoder by training on the full training set and predicting target directions on the test set. For comparison, we also decoded from firing rates using the same procedure. Firing rates were computed as described above in *Decoding of movement kinematics*.

To compute performance of the position decoder, we predicted the x- and y-position in the relevant reference frame, then computed the general coefficient of determination (r^2^) between the true and predicted values on each held-out set. To obtain a single performance score, we averaged the r^2^ for the x- and y-coordinate, weighted by the variance in each dimension. For the direction and magnitude decoders, we first normalized each target position vector to the unit circle. We then jointly predicted the x- and y-coordinate of the normalized vector along with the magnitude of the unnormalized vector. The performance of the direction decoder was defined as the average of the r^2^ between the predicted and actual x- and y-coordinates. Cartesian coordinates were chosen instead of angles to eliminate wraparound issues for decoding.

The best-performing hyperparameters on the hyperparameter training set were used to evaluate the model using the test set, and to compare hand-centric and shoulder-centric decoding using cross-validation. Generalization performance was evaluated using randomly shuffled 10-fold cross validation using the test set. This was repeated 10 times, for a total of 100 splits of the data. Both the hand-centric model and the shoulder-centric model were fit and tested using each of the 100 splits to get an estimate of the generalization performance of the models. To test the significance of the difference between the hand-centric and shoulder-centric model, we applied a paired-sample t-test with corrected variance for cross-validation^63^.

### Comparing coding of initial and corrective submovements

We binned the angle of the target direction relative to the hand into 12 equal-sized bins. For each bin, we computed the mean value of inferred input in a window from 350 to 50 ms before the initial movement. We performed the same procedure for corrective submovements, except using the same window relative to the corrective submovement.

This resulted in 24 direction-means each for the initial submovements and the corrective submovements (12 direction bins × 2 inferred inputs). We computed the Pearson correlation coefficient between the direction-means for the initial submovements and the corrective submovements. To combine across the two inferred inputs when computing this correlation, we first z-scored the direction-means in each movement condition before computing the correlation coefficient.

For Figure 7C, we performed this same procedure, except that instead of computing the means of inferred input for each direction bin, we computed the mean of the first 5 principal components of the smoothed firing rates (see above). This was done to denoise the tuning curve given that there were fewer corrective submovements than initial movements.

### Simulations

We simulated a continuous-rate recurrent neural network (RNN) which controlled the joint angle velocities of a two-link arm completing the random target pursuit task. Each simulated trial required movements of the “hand” to a series of two targets. The network consisted of three layers: an input layer, a recurrent layer, and an output layer. The input layer consisted of two units which received the position of the current target in x and y, respectively. The encoding of the target by the input layer varied by condition, as described in the next section. The activation of the recurrent layer with *N* units at time *t* followed the discrete time equation:

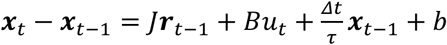

where 𝒙_t_ ∈ ℝ^N^ is the vector of activations at time *t*, 𝐽 ∈ ℝ^N×N^ is the matrix of recurrent weights, 𝑢_t_ ∈ ℝ^2^ are the inputs at time *t*, 𝐵 ∈ ℝ^N×2^ is the matrix of input weights, *b* is a trainable bias term, and *𝜏* is the time constant of the activation. The recurrent rates are related to the activations as follows:

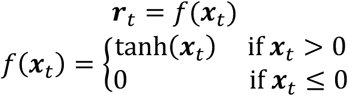

The outputs 𝒛_t_ ∈ ℝ^2^ were determined by a linear readout of trainable weights:

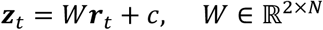

The two outputs represent the joint velocities of the two joints in the arm. For all simulations, we used *N* = 100 units, Δ*t* = 10 ms, and *𝜏* = 50 ms.

### Inputs

The inputs represent the target according to one of four conditions: sustained input in shoulder-centered coordinates, sustained input in hand-centered coordinates, transient input in shoulder-centered coordinates, and transient input in hand-centered coordinates. The sequence of inputs starts with a zero pad lasting 50 ms. Onset jitter was introduced by including a sequence of zeros with duration drawn randomly from a uniform distribution ranging from 0 to 60 ms. After the zero pad and jitter, the inputs with values corresponding to the location of the target began. For the sustained input conditions, the informative inputs continued until the next input. For the transient inputs, the informative inputs lasted one time step before returning to zero. The input specifying the second target occurred after the first reach was completed, plus another 0-60 ms jitter interval.

### Outputs

The model was trained to produce joint velocities which corresponded to reach trajectories that acquired each target. The reach trajectories were straight lines from the initial position of the end effector to the target with a Gaussian speed profile. The speed profile of each reach had a s.d. of 20 ms and extended for 4 * s.d. in each direction, for a total of 160 ms per reach.

To produce the training output, we generated two random target locations from a uniform distribution over the 2D workspace. Then a reach trajectory in Cartesian coordinates was generated corresponding to a straight line between the starting position of the end effector and the target with a Gaussian speed profile. The trajectory was converted into a sequence of joint velocities by solving the inverse kinematics problem as follows.

Let L_1_ and L_2_ be the fixed lengths of the first and second arm components, respectively, and let 𝜃_1_, 𝜃_2_ be the time-varying angles of the joints, and 𝒙 = (𝑥_1_, 𝑥_2_) ∈ ℝ^2^ be the position of the end effector. Then

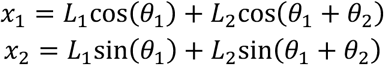

From these equations we can compute the Jacobian matrix J = d***x***/d𝜃. From the chain rule, we have ***x****′* = J 𝜃***′***, meaning 𝜃*′* = J^-1^ ***x****′*. This allows us to compute the sequence of joint velocities in terms of the sequence of end effector velocities and positions in Cartesian coordinates.

### Training

The network was trained to minimize the mean squared error of the outputs with respect to the target sequence of joint angle outputs. We applied three regularization terms to the objective. The first was a standard weight regularization, penalizing the 2-norm of weights in the input and output layers of the network. The second regularization was a penalty on the square of the rates in the recurrent layer at every time step. This prevents rates from saturating at their maximal value. The third regularization was a penalty on the smooth maximum over the units of the activity on the final timestep of the trial:

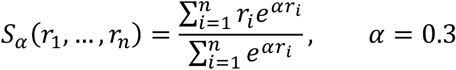

with *r_i_* the final firing rate of the *i*^th^ neuron.This helps create naturalistic solutions with activity that peaks during the reach, rather than moving from one nonzero fixed-point attractor to another.

The network was trained using the ADAM optimizer (Kingma and Ba 2014). We trained one network for each condition, using a training set of 2048 randomly generated trials with two targets each. The fitted networks were then simulated on 512 validation trials with targets unseen during training. The activity on the validation trials was fit using LFADS. In order to pass the simulation data to LFADS, we converted the continuous activity of the network to spike activity by simulating an inhomogeneous Poisson process for each trial and neuron, using a maximum instantaneous rate of 100 spikes/s.

### LFADS ensembles

The ensembles of models for both the simulated and monkey data analysis were constructed by varying two hyperparameters: the cost of correlation (through time) in the means of controller output (c_co_mean_corr_scale) and the initial autocorrelation of AR(1) priors (in time bins) (c_prior_ar_atau). When varying the latter hyperparameter, another hyperparameter corresponding to training the autocorrelation length (c_do_train_prior_ar_atau) was also set to False. The rest of the training procedure was left untouched after initially varying these hyperparameters during LFADS model specification.

### Hyperparameters

Increasing the c_co_mean_corr_scale hyperparameter from zero decorrelates the output, which we speculated would allow for more transient types of peaks in the controller output (Fig. 5b, middle column).

Modifying the c_prior_ar_atau hyperparameter allowed us some influence on the timescale of the controller fluctuations. By not allowing the model to update this autocorrelation length (switching the hyperparameter c_do_train_prior_ar_atau to False) this timing specification of the controller outputs persisted throughout training.

### Simulated Data

For the simulations, we tested two input timing schemes and three hyperparameter combinations per timing scheme. For both the transient and sustained timing schemes, the base LFADS hyperparameter sets that we used on the monkey data (Table 1) constituted the partial-match cases.

For both timing schemes we set the c_co_mean_corr_scale hyperparameter to 0.7 (Table 2), which for the transient inputs corresponded to the match case and for the sustained inputs corresponded to the mismatch case.

We set the c_prior_ar_atau hyperparameter very short (2 bins) in addition to decorrelating the outputs for the sustained-mismatch case. We lengthened this hyperparameter (30 bins) for the transient-mismatch case and for the sustained-match case (Table 2).

### Monkey Data

For the monkey data, we varied the two hyperparameters to form three additional LFADS models for the ensemble along with the original model (Original, Fig. 3 and Table 1; Additional, Fig. 6 and Table 3).

Both autocorrelation models were trained with c_do_train_prior_ar_atau set to False (as for the simulations) and the Short Autocorrelation model was trained with c_prior_ar_atau set to 2 bins width, while the Long Autocorrelation model had c_prior_ar_atau set to 30 bins width. The Decorrelated model was trained with c_co_mean_corr_scale set to 0.7.

## Acknowledgements

The authors thank Josh Coles, Zach Haga, Jignesh Joshi, Dawn Paulsen, Jacob Reimer, Aaron Suminski, and Dennis Tkach for originally collecting the data, and Chethan Pandarinath for assistance with LFADS. This work was funded by R01 NS111982 (NGH), R01NS045853 (NGH) R01 EY022338 (JNM), NSF CAREER Grant 0952686 (JNM), R01-NS094184 (JNM), The Whitehall Foundation (MTK), The Simons Collaboration on the Global Brain (MTK), NIH R01 NS121535 (MTK), the NSF-Simons Institute for Theory and Mathematics in Biology (MTK), The University of Chicago (MTK), the Neuroscience Institute at the University of Chicago (MTK), and T90/R90 T90DA059109 (AV).

---

**Supplementary Figure 1.**
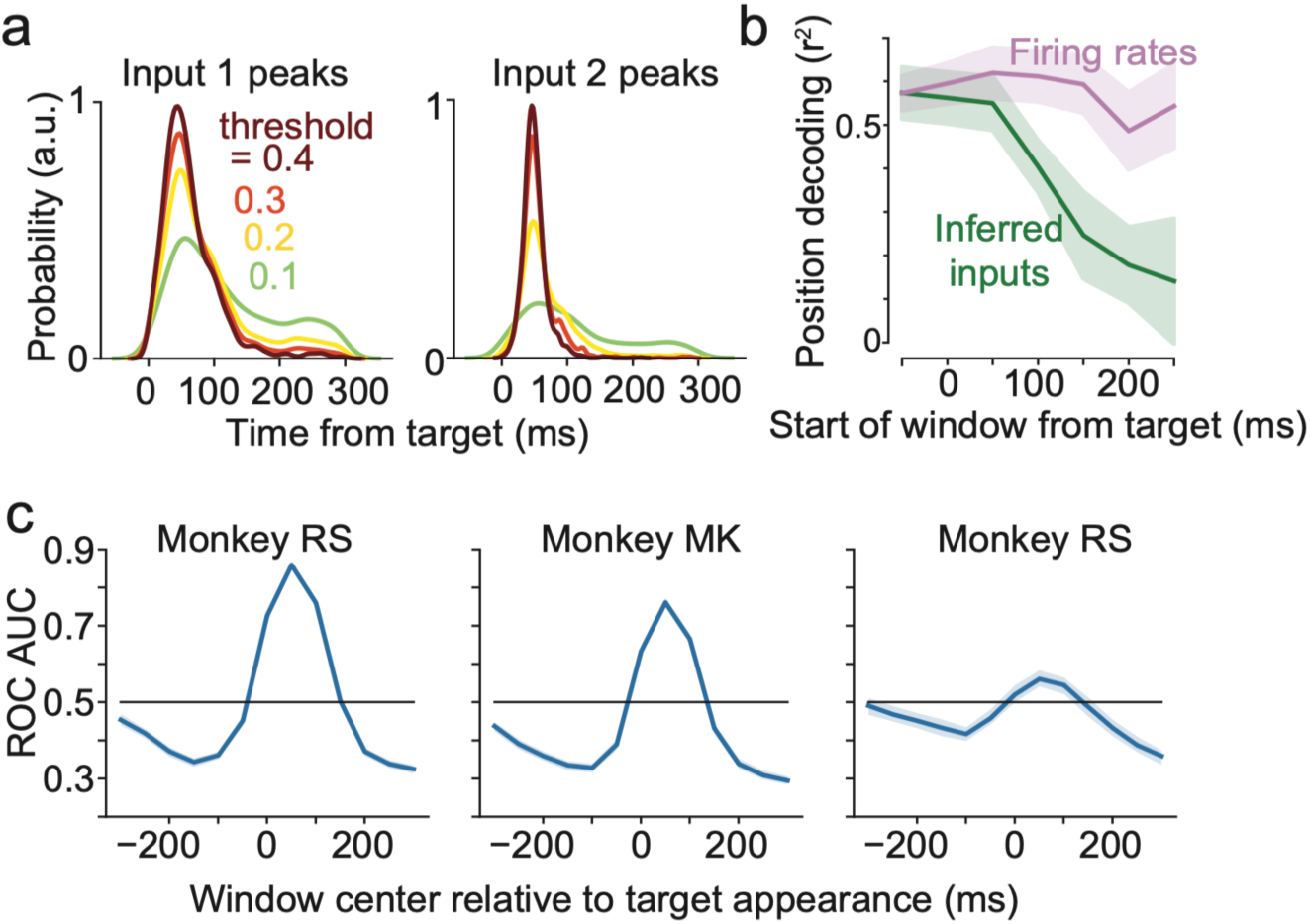
Additional comparisons for inferred initial-movement inputs on monkey data. **(a)** Distribution of the timing of peaks in the inferred input relative to the appearance of a new target. Distributions are shown for four different threshold values above which peaks are included. Distributions for monkey RS. **(b)** Cross-validated decoding performance of target position prediction using different time windows. X-axis values are centers of 250 ms windows used to predict target direction. **(c)** Area under the receiver operating characteristic (ROC) curve showing how reliably particular windows in time can be distinguished from the rest of the trial using controller magnitude. Values for a range of 100 ms windows (see Methods). Shaded error is 95% bootstrapped confidence interval (the error region at some points may be narrower than the line thickness).

**Supplementary Figure 2.**
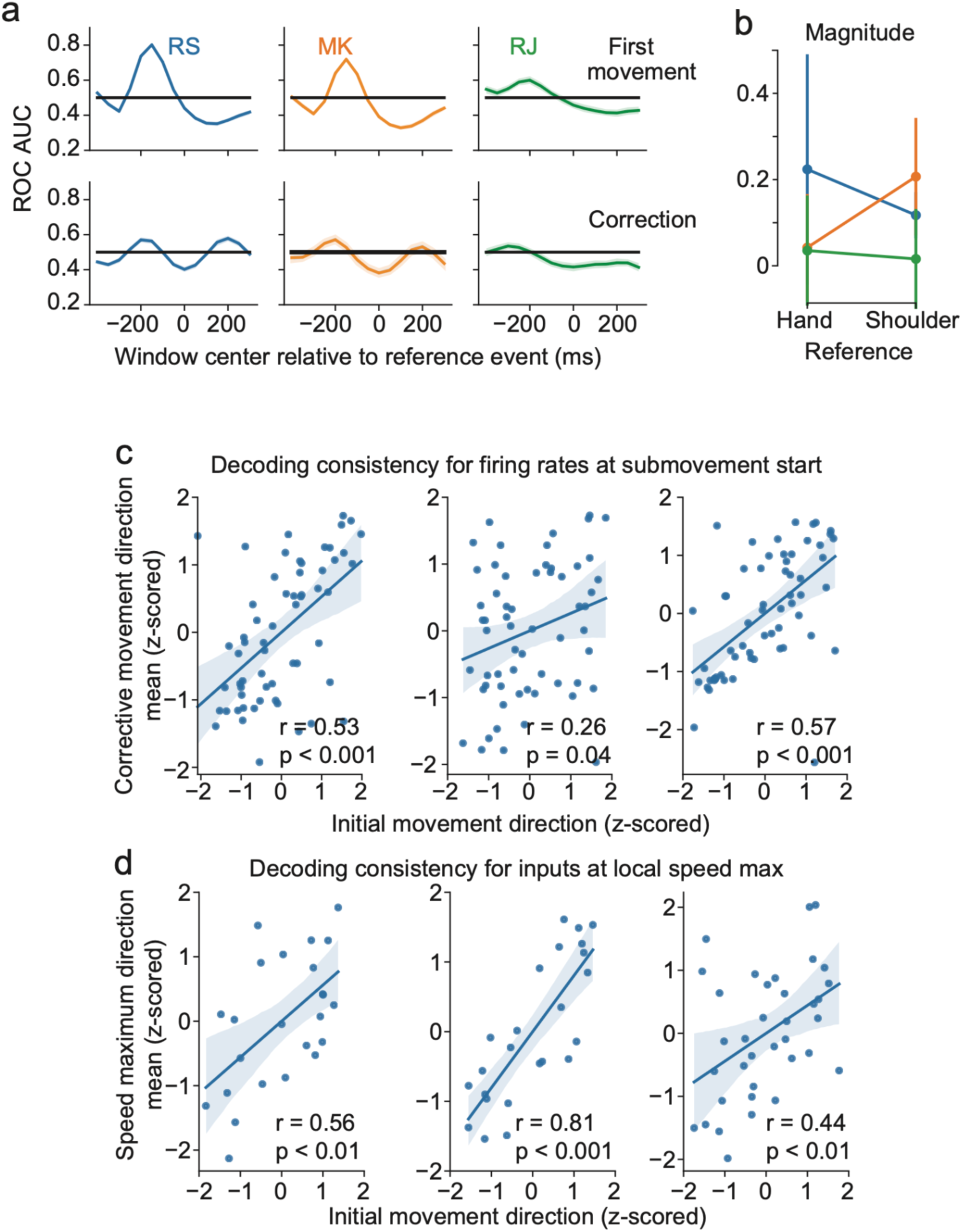
Additional comparisons for inferred corrective-movement inputs on monkey data. **(a)** Area under ROC curve across different windows as in Supplementary Figure 1c, but relative to movement onset for initial and corrective movements rather than target appearance. Shaded error is 95% bootstrapped confidence interval (the error region at some points may be narrower than the line thickness). **(b)** Magnitude decoder for the same data as in Figure 4d,e. **(c)** Same as Figure 4g but using 5 principal components of firing rates rather than controller values. **(d)** The correlation of direction-averaged inferred input around the initial movement to direction-averaged input around the maximum speed.

**Supplementary Table 1.**
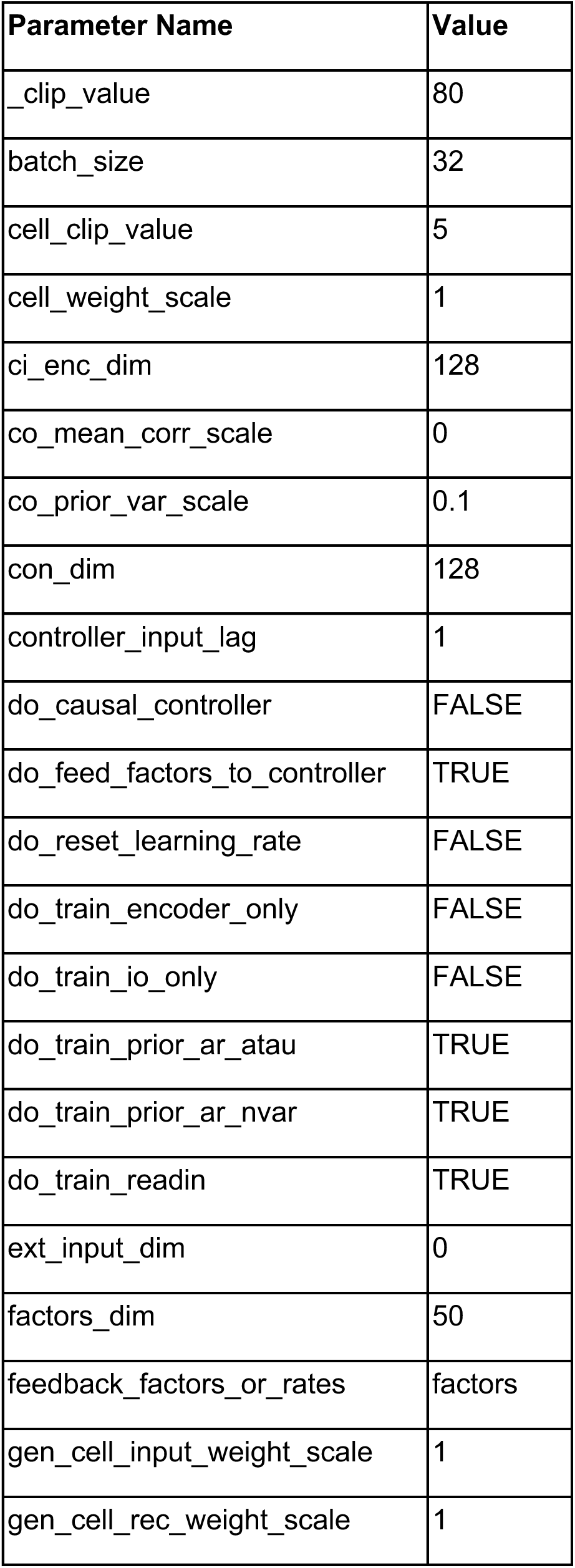
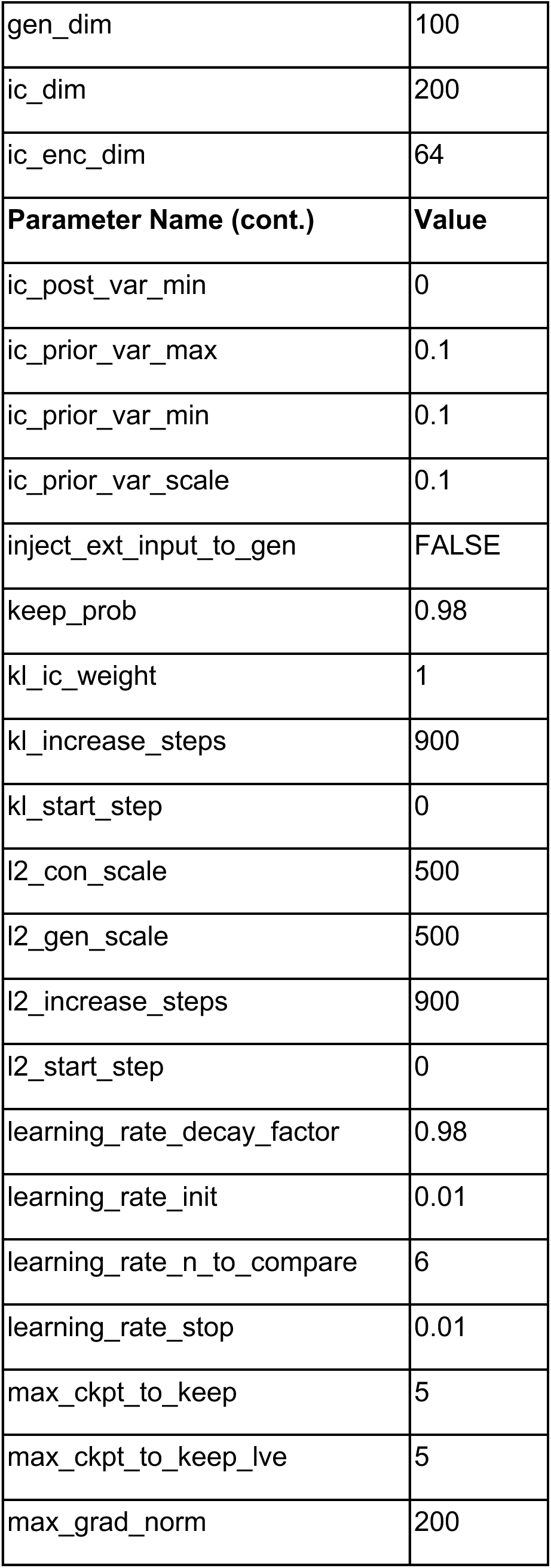
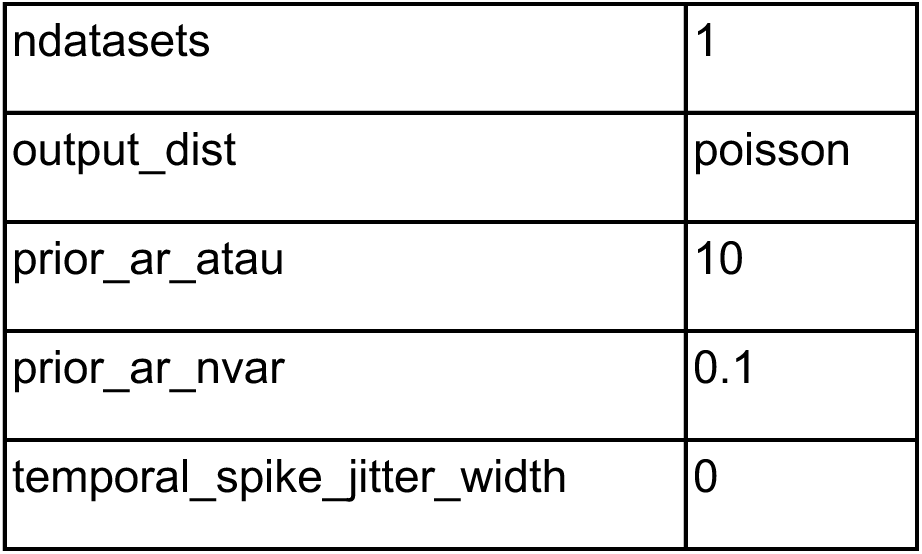
Hyperparameter values for fitting LFADS. Parameter names are as in the code package.

| Parameter Name | Value |
| --- | --- |
| _clip_value | 80 |
| batch_size | 32 |
| cell_clip_value | 5 |
| cell_weight_scale | 1 |
| ci_enc_dim | 128 |
| co_mean_corr_scale | 0 |
| co_prior_var_scale | 0.1 |
| con_dim | 128 |
| controller_input_lag | 1 |
| do_causal_controller | FALSE |
| do_feed_factors_to_controller | TRUE |
| do_reset_learning_rate | FALSE |
| do_train_encoder_only | FALSE |
| do_train_io_only | FALSE |
| do_train_prior_ar_atau | TRUE |
| do_train_prior_ar_nvar | TRUE |
| do_train_readin | TRUE |
| ext_input_dim | 0 |
| factors_dim | 50 |
| feedback_factors_or_rates | factors |
| gen_cell_input_weight_scale | 1 |
| gen_cell_rec_weight_scale | 1 |
| gen_dim | 100 |
| ic_dim | 200 |
| ic_enc_dim | 64 |
| **Parameter Name (cont.)** | **Value** |
| ic_post_var_min | 0 |
| ic_prior_var_max | 0.1 |
| ic_prior_var_min | 0.1 |
| ic_prior_var_scale | 0.1 |
| inject_ext_input_to_gen | FALSE |
| keep_prob | 0.98 |
| kl_ic_weight | 1 |
| kl_increase_steps | 900 |
| kl_start_step | 0 |
| l2_con_scale | 500 |
| l2_gen_scale | 500 |
| l2_increase_steps | 900 |
| l2_start_step | 0 |
| learning_rate_decay_factor | 0.98 |
| learning_rate_init | 0.01 |
| learning_rate_n_to_compare | 6 |
| learning_rate_stop | 0.01 |
| max_ckpt_to_keep | 5 |
| max_ckpt_to_keep_lve | 5 |
| max_grad_norm | 200 |
| ndatasets | 1 |
| output_dist | poisson |
| prior_ar_atau | 10 |
| prior_ar_nvar | 0.1 |
| temporal_spike_jitter_width | 0 |

**Supplementary Table 2.** Hyperparameter value changes for LFADS ensemble models on simulated data. Parameter names are as in the code package.

| LFADS Model on Simulated Data | Parameters Changed (from Table 1) | Value |
| --- | --- | --- |
| Transient-match | c_co_mean_corr_scale | 0.7 |
| Transient-partial-match | None | N/A |
| Transient-mismatch | c_prior_ar_atau | 30 |
|  | c_do_train_prior_ar_atau | False |
| Sustained-match | c_prior_ar_atau | 30 |
|  | c_do_train_prior_ar_atau | False |
| Sustained-partial-match | None | N/A |
| Sustained-mismatch | c_prior_ar_atau | 2 |
|  | c_do_train_prior_ar_atau | False |
|  | c_co_mean_corr_scale | 0.7 |

**Supplementary Table 3.** Hyperparameter value changes for ensemble LFADS models on monkey data. Parameter names are as in the code package.

| LFADS Ensemble Model | Parameters Changed (from Table 1) | Value |
| --- | --- | --- |
| Short Autocorrelation | c_prior_ar_atau | 2 |
|  | c_do_train_prior_ar_atau | False |
| Long Autocorrelation | c_prior_ar_atau | 30 |
|  | c_do_train_prior_ar_atau | False |
| Decorrelated | c_co_mean_corr_scale | 0.7 |

